# Integrating process, proximity, and prediction implicates novel protein and RNA interactions in human Origin Recognition Complex function

**DOI:** 10.1101/2025.05.28.651956

**Authors:** John A. Smolka

## Abstract

Genome replication start sites, called origins, begin to be specified by Origin Recognition Complex (ORC) proteins prior to replication through a process called origin licensing. Once licensed, origins become active and initiate DNA synthesis with varying efficiencies influenced by local chromatin environment, transcription, and 3D genome organization. ORC proteins have also been implicated in regulating chromatin state and nuclear organization. However, it is unclear if there is interplay between ORC and the chromatin architectures underlying origin activation, as we lack a systems-level understanding of how ORC proteins interact, post-licensing, with the nuclear environments conducive to genome synthesis. To infer this context-specific ORC interactome, I used data from genome-wide CRISPR fitness screens, a cell-wide proximity labeling study, and proteomic profiling of nascent DNA-associated proteins to identify 17 novel factors that genetically and proteomically interact with ORC subunits and genome replication. Unexpectedly, the candidate pool was significantly enriched for factors involved in the homeostasis of RNA Polymerase III (Pol III) transcripts, particularly 5S ribosomal RNA (rRNA) and transfer RNA (tRNA). Follow-up protein-protein structure predictions by AlphaFold 3 (AF3) proposed direct interactions between ORC subunits and Pol III transcript biogenesis factors, as well as epigenetic regulators and a cyclin-dependent kinase. Given the prominence of 5S rRNA and tRNA biogenesis factors in my results, and prior reports of ORC subunits binding RNA, I also used protein-RNA structure prediction to identify candidate ORC3 interactions with 5S rRNA and tRNA. Altogether, my analysis integrates biological process, molecular proximity in human cells, and structural prediction to nominate novel protein and RNA interactions for involvement in the human replication program. These results augment and expand current models for ORC function and origin activation, particularly those involving chromatin state and transcriptional activity, and generate testable hypotheses to explore the interdependencies of replication patterning, histone modification, and nuclear RNA homeostasis.

## Introduction

Organisms must duplicate genomic DNA to grow, replenish tissues, and reproduce. Proliferating human cells, typically, must complete one round of DNA replication during S phase of the cell cycle to provide intact genetic information for two daughter cells. Because the diploid human genome is approximately 6 billion basepairs in size, DNA synthesis must coordinately initiate from thousands of genomic sites, termed origins of replication, to complete replication within the span of S phase.^1^ To meet this need, human cells require an ensemble of proteins that coordinates DNA synthesis in time and space to efficiently complete one round of replication before division. Defects in the timing and patterning of replication initiation, also called the replication program, can compromise genome integrity and contribute to genetic disease and cancer.^2,3^ Accordingly, the replication program undergoes characteristic changes throughout cellular differentiation and also becomes abnormal in cancer cells.^4,5^ These observations emphasize the need for complete mechanistic models of the replication program in normal and pathological contexts to more effectively engineer cell fate and therapeutically target cancer vulnerabilities. However, the molecular determinants of when and where human genome replication initiates are still being characterized.

One of the most well-established effectors of replication patterning in Eukaryotes is the Origin Recognition Complex (ORC).^1^ ORC is a DNA-binding protein assembly, consisting of 6 subunits (ORC1-6), that establishes origins of replication through sequential cell cycle-timed events. ORC subunits form a chromatin-associated holocomplex as early as M phase and throughout G1.^6^ During G1, ORC specifies origins through a process called origin licensing, when ORC subunits assemble in an ATP-dependent manner into stable DNA-bound complexes with the ATPase CDC6; this binding appears to be DNA sequence-independent, and guided primarily by chromatin accessibility.^7^ Then, through interactions with ORC and the helicase loader CDT1, the MCM replicative helicase is recruited to ORC-bound sites, creating licensed origins with resident pre-replicative complexes (pre-RCs).^1,4^ Pre-RCs poise licensed origins for the formation of a pre-initiation complex (pre-IC), assembly of a complete replisome, and, ultimately, the initiation of DNA synthesis in S phase. In addition to their roles in licensing, some ORC subunits have also been implicated in heterochromatin maintenance and nuclear organization, but these roles are currently thought to be independent of ORC’s function in the replication program.^8,9^

Though critical for establishing origin identity, origin licensing does not commit an origin to replication initiation. Licensed origins are typically in excess, and only a subset becomes active while many remain dormant.^1^ Furthermore, origins activate at different times throughout S phase, being classified generally as “early” and “late”. While ORC-mediated origin licensing in M and G1 phases is mechanistically well-understood, the processes downstream of licensing that lead to selective and timed origin activation in S phase, and ORC’s participation in those processes, remain unclear. While some studies suggest that ORC is removed from chromatin after licensing completes^10^, other studies propose that ORC components remain chromatin-bound, possibly even at origins to participate in origin activation.^11–13^ Regardless, evidence supports that, after licensing, suppression of re-replication is accomplished by cyclin-dependent kinase-mediated phosphorylation of ORC subunits and the degradation of ORC1, both of which inhibit licensing.^6,14^ Taken together, these data suggest that some ORC subunits remain chromatin-associated through the G1-S transition in a licensing-inhibited state, but it is unclear if they remain in the molecular environment of origins and/or effect additional regulation of the replication program.

While roles for ORC in origin activation are debatable, the chromatin environments that correlate positively with origin activation are increasingly well-characterized. Data indicate that replication initiation is probabilistic, with some regions of the genome being “high efficiency” initiation sites that tend to be early origins.^15^ Origin mapping by a variety of methods has consistently identified early origins in intergenic regions proximal to active gene promoters, suggesting origin activity correlates with open chromatin, local transcriptional activity, and histone acetylation, which promotes all of the above.^16,17^ It is thought that histone acetylation creates open chromatin environments permissive for ORC binding and origin licensing.^7^ Once pre-RCs have formed, chromatin acetylation can promote replication initiation independently of ORC through recruitment of bromodomain-containing proteins BRD2 and BRD4, which in turn can recruit the replication initiation factor TICRR, also known as Treslin, that facilitates the transition of pre-RCs to pre-ICs.^18,19^ Accordingly, targeting histone acetyltransferases (HATs) and deacetylases to replication origins has been shown to advance and delay replication timing, respectively.^20^ Additionally, a study in yeast found that histone acetylation at origins is dynamic, increasing at the G1-S transition and peaking in S phase and G2.^21^ This suggests that chromatin acetylation not only predisposes loci to becoming licensed and active origins, but also undergoes amplification at origins during the G1-S transition and S phase. In line with this, the histone acetyltransferase HBO1 has been identified as an interactor of CDT1, and thus a tertiary ORC interactor, that acetylates histones at origins during G1.^22,23^ Altogether, this body of knowledge indicates ORC and histone acetylation act in concert to shape the replication program, and hint at ORC- and acetylation-dependent feedback mechanisms influencing origin specification and activation.

From chromatin state to transcription and 3D genome organization, a confluence of nuclear processes has been linked to both ORC function and origin activation.^1,4,15,24,25^ However, cause-and-effect relationships between an origin’s licensing, activation, and local nuclear environment remain elusive. This stems in part from limited knowledge of ORC’s molecular interactions with the nuclear milieu, particularly at and after the G1-S transition. This gap in knowledge precludes a systems-level understanding of ORC function that could help clarify the molecular processes influencing the probability that an origin will become active once licensed. To explore these possibilities, I first used a dynamic modeling approach to demonstrate that a chromatin-based model for origin activation is insufficient without positive feedback, and that ORC is an ideal conduit for this feedback. Guided by this model, I hypothesized the human replication program relies on an ensemble of yet-to-be-defined molecular interactions with ORC proteins post-licensing and during S phase. To probe this hypothesis, I integrated publicly available CRISPR fitness screen and cellular proximity labeling data to derive a dual-criteria ORC interactome constituted by both genetic and proteomic interactions, and used nascent DNA proteomics data to infer the portion of this interactome overlapping the protein environments of active DNA replication. This refined ORC interactome contained 17 novel candidates, many involved in noncoding RNA (ncRNA) metabolism, that have not been previously described as having roles in genome maintenance. I then used multimeric structure prediction to estimate the likelihood that ORC subunits form complexes with candidate proteins and relevant small ncRNA species. These predictions posited an ensemble of novel protein and RNA interactions linking ORC function to RNA biogenesis and processing, epigenomic processes, and a cyclin-dependent kinase. The proposed interactions synergize with my dynamic model for origin activation to generate new hypotheses about ORC protein participation in the replication program beyond origin licensing.

## Methods

### Modeling and simulation of origin activation

Logical models were built and model outputs were simulated using the Cell Collective platform (https://cellcollective.org/). Simulations were run with synchronous updating and default parameters, with the “acetyltransferase” external input set at 20% activity, which was empirically determined to be the optimal activity level for observing a dynamic range of model outputs within 100 steps. All logical model .sbml files can be made available upon request.

### Acquisition and analysis of genetic and proteomic screen data

2Q24 CRISPR gene effect data was downloaded via the DepMap online portal (https://depmap.org/portal/). Proximity labeling/BioID data (SAINTexpress report) was downloaded via the Human Cell Map online portal (https://cell-map.org/); datasets were formatted to be gene x cell line and protein x bait matrices, respectively. DepMap and Human Cell Map datasets were analyzed using R programming language in RStudio. Pairwise correlations were derived by the corr() function using “pairwise.complete.obs”. Hierarchical clustering, cluster annotation, and heatmap generation were performed using correlation matrices and heatmap() and heatmap.2() functions.

### Network and gene ontology analysis of gene/protein ensembles

Human gene/protein lists were queried for reported interactions and gene ontology enrichment using the STRING database (https://string-db.org/). Interaction network representations were generated using experiments and databases as active interaction sources.

### Biomolecular structure prediction and visualization

Structural predictions were performed using AlphaFold 3 via the AlphaFold server (https://golgi.sandbox.google.com). Three randomly seeded (default setting) predictions were run per ORC subunit-candidate protein pair, and the average ipTM value was used for initial screening. Six additional randomly seeded predictions were performed for protein pairs that met cut-off criteria; analyses were performed identically for protein-RNA pairs. The .model files from AlphaFold 3 outputs for select predicted structures were three-dimensionally rendered as molecular surface models and posed using the Mol* 3D Viewer tool provided by the RCSB Protein Data Bank (https://www.rcsb.org/3d-view). All AlphaFold 3 outputs for structures presented can be made available upon request.

### Statistics

P-values were determined without assumption of normality using the wilcox.test function in R. Significance is represented as follows: * for p ≤ 0.05, ** for p ≤ 0.01, *** for p ≤ 0.001, and **** for p ≤ 0.0001. Statistical outliers were defined as ≥ 2 standard deviations from the mean.

## Results and discussion

### A minimal dynamic model for origin activation motivates exploration of ORC interactions downstream of origin licensing

Current molecular models for replication initiation are linear, treat chromatin state as static, restrict ORC’s activities to licensing, and/or do not explain the probabilistic nature of origin activation. Thus, to provide a conceptual framework for probing new hypotheses about ORC function and origin activation, I used the Cell Collective platform to create a nonlinear, dynamic network model of origin activation integrating chromatin state, ORC, and origin licensing.^26^ Studies of origin licensing and activation indicate that chromatin acetylation promotes both processes. I reasoned that these relationships constitute a feedforward network^27^, where i) histone acetylation promotes open chromatin, open chromatin facilitates ORC association with DNA, which then permits origin licensing and pre-RC formation, and ii) histone acetylation, in parallel, recruits acetylation readers like BRD2/4, which recruit TICRR, which promotes the formation of pre-ICs if pre-RCs are pre-existing [Fig.1A]. Thus, in this model, initiation is dependent on origin licensing by ORC, but can also be regulated independently of ORC. I further reasoned that origin activation can be treated as a bistable process, in which replication initiation is a single, all-or-nothing event that irreversibly transitions an origin from inactive to active; similar to a cell cycle transition, this event is thresholded, requiring a confluence of events to transduce a variable input into a switch-like output.^28^ I tested these assumptions using a Boolean logical model with a single input, acetyltransferase activity, and a single output, origin activation [Fig.1A]. While this simplified model does not take into account cell cycle progression, it incorporates a conditional requiring the pre-RC node be active for positive regulation of the pre-IC node to occur, emulating the sequential nature and directional dependency of origin licensing and activation.

**Figure 1.**
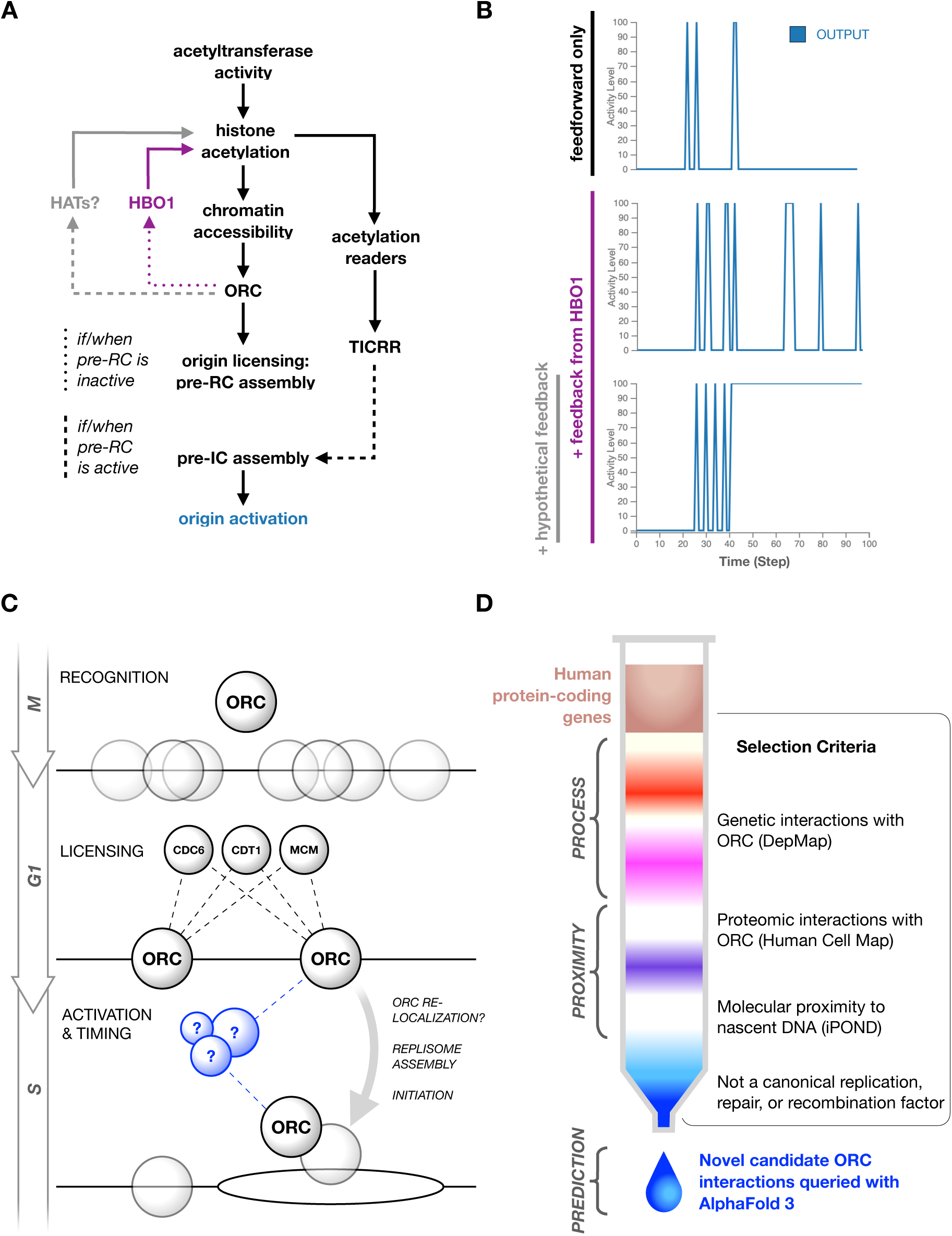
(A) Network model for origin activation integrating chromatin state and ORC function. (B) Representative outputs over time from simulations testing iterations of the model in A as Boolean logical models with: no positive feedback (feedforward only), positive feedback representing an ORC-mediated interaction with HBO1 before licensing (+ feedback from HBO1), and additional positive feedback representing hypothetical ORC-mediated interactions with HATs after licensing (+ hypothetical feedback). (C) Simplified schematic for replication patterning with ORC-mediated origin licensing and hypothetical ORC interactions after licensing. (D) Schematic of a data science pipeline for isolating and identifying candidate ORC interactions of interest from large datasets, analogized to chromatography.

Plots of simulated output over time from this feedforward network produced qualitatively brief and stochastic “on/off” oscillations (“on” being 100% activity, “off” being 0%) [Fig.1B], which I interpret as output instability and activation failure. These observations demonstrate this positive feedforward regulation alone is not sufficient to create a bistable output under the conditions used. Bistability can be promoted by feedback mechanisms.^29,30^ Given this premise, I wondered if introducing positive feedback on histone acetylation from a downstream component could create a bistable output. Based on the documented functional relationship between HBO1 and licensing^22^, I hypothesized that positive feedback from ORC on histone acetylation through HBO1 could introduce bistability to the output of this network. Since HBO1 activity is inhibited after licensing/G1^23^, I used a conditional preventing HBO1 activity when the pre-RC node becomes active, emulating the restriction of HBO1 activity to the licensing phase or earlier. This modified logical model produced significantly more frequent on/off oscillations (mean number of oscillations within 100 steps = 2.8, baseline, versus 7.1 with HBO1-mediated feedback added, p = 2.56e-6), but also failed to achieve bistability under the conditions used [Fig.1A,B,S1A]. Since introducing constrained positive feedback enhanced activity oscillations, I wondered if introducing additional feedback unconstrained by pre-RC activity would further enhance this trend or achieve bistability. To test this, I introduced a duplicate, hypothetical HAT-mediated positive feedback loop from ORC onto histone acetylation with a conditional permitting hypothetical HAT activity only when the pre-RC node is active. This iteration of the logical model exhibited bistable behavior distinct from the other two models, producing switch-like patterns of output with variable periods of steady-state oscillations followed, in all cases, by a permanently active state [Fig.1A,B]; I note there were rare events with immediate entry into a permanent active state, with no oscillations prior [Fig.S1B]. Although simplified, this model proposes ORC-mediated feedback that amplifies histone acetylation around origins, both before and after pre-RC assembly, as a way to promote switch-like, bistable origin activity.

Broadly, these results inform a hypothesis that ORC interactions acting at the transition into and during S phase could provide ideal vectors for promoting replication initiation. Though ORC becomes resident at numerous genomic sites through origin recognition and licensing in M and G1 phases, ORC’s molecular interactions and relationship with origin activity as G1 transitions into S phase remain unclear [Fig.1C]. To probe for ORC interactions associated with active origins, I chose three high-throughput datasets to infer genetic, subcellular, and replication-specific ORC interactomes. To infer functional dependencies, I leveraged genetic interaction data from the Cancer Dependency Map Project (DepMap). The DepMap consortium has performed genome-wide CRISPR fitness screens in over 1,000 different cancer cell lines.^31^ Growth effect correlations, or co-essentialities, derived from these data have been used to map genetic pathways and identify novel factors in processes of interest.^32,33^ To infer subcellular interactions, I utilized proteomic data from the Human Cell Map.^34^ This dataset originated from a proximity labeling study that employed BioID with 192 subcellular markers to spatially map the human proteome at subcompartmental resolution in cells. I confirmed both DepMap and Human Cell Map datasets identify expected genetic and proteomic interactions across the core human replication machinery, including ORC subunits, and quantitatively distinguish these interactions from general nuclear protein interactions [Fig.S2A-D], supporting the suitability of these datasets for this study. Finally, I referenced these interactomes with the proteome identified by the isolation of proteins on nascent DNA, or iPOND.^35^ This methodology has enabled the proteomic inference of proteins physically associated with the genome replication process and, excitingly, found that ORC subunits 2-6 are enriched in these protein environments. I thus designed an analytical pipeline using each of these datasets as selective criteria to identify novel candidate ORC interactions with the potential to occur in the environments of active replication [Fig.1D]. This pipeline nominates candidates first by data to then be screened for the likelihood of protein-protein interactions with ORC subunits using multimeric structure prediction by AlphaFold.^36^ I term this approach an integration of process (DepMap), proximity (Human Cell Map and iPOND), and prediction (AlphaFold).

### Integration of data from CRISPR fitness screens and proteomic studies infers a novel network of protein interactions supporting ORC function and genome replication

Using biological process as my first criterion, I identified a subpopulation of human protein-coding genes that genetically interact with ORC. I derived the ORC genetic interactome by hierarchically clustering all protein-coding genes currently available in DepMap data by growth effect correlations with ORC subunits 1-6 [Fig.2A]. This identified a cluster of 785 genes enriched for positive growth effect correlations with ORC subunits, with a significantly higher mean Pearson correlation (^r̅^) than the mean ORC subunit correlation with all genes (^r̅^ = 0.0046 for all, versus ^r̅^ = 0.0971 for cluster, p < 2.22e-308) [Fig.2B]. Expectedly, this cluster contained 81 established genome maintenance factors, including multiple DNA repair proteins as well as subunits of the replication clamp-loading RFC complex, leading and lagging strand polymerases, MCM replicative helicase, and GINS complex that promotes replisome assembly and function [Fig.S3]. Notably, ORC subunits 2-5 were included in this cluster, but primary effectors of origin licensing ORC1, CDC6, and CDT1 were not, suggesting that the co-essentialities identified do not reflect the origin licensing process occurring in M and G1 phases. Unexpectedly, the most enriched biological process in this cluster was mitochondrial gene expression, followed by noncoding/translational RNA metabolism [Fig.2C]. Though unexpected, a recent synthetic lethality screen in human cells also found mitochondrial gene expression to be the most enriched term amongst factors identified as essential for replication stress survival^37^; RNA homeostatic processes have also been identified in genetic screens for factors required for genome stability.^38^ Altogether, this suggests that the network of ORC co-essentialities identified by my analysis reflects a combination of direct players in genome maintenance and replication, cellular pathways that combat replication stress and support genome stability, and facets of ORC function beyond origin licensing.

**Figure 2.**
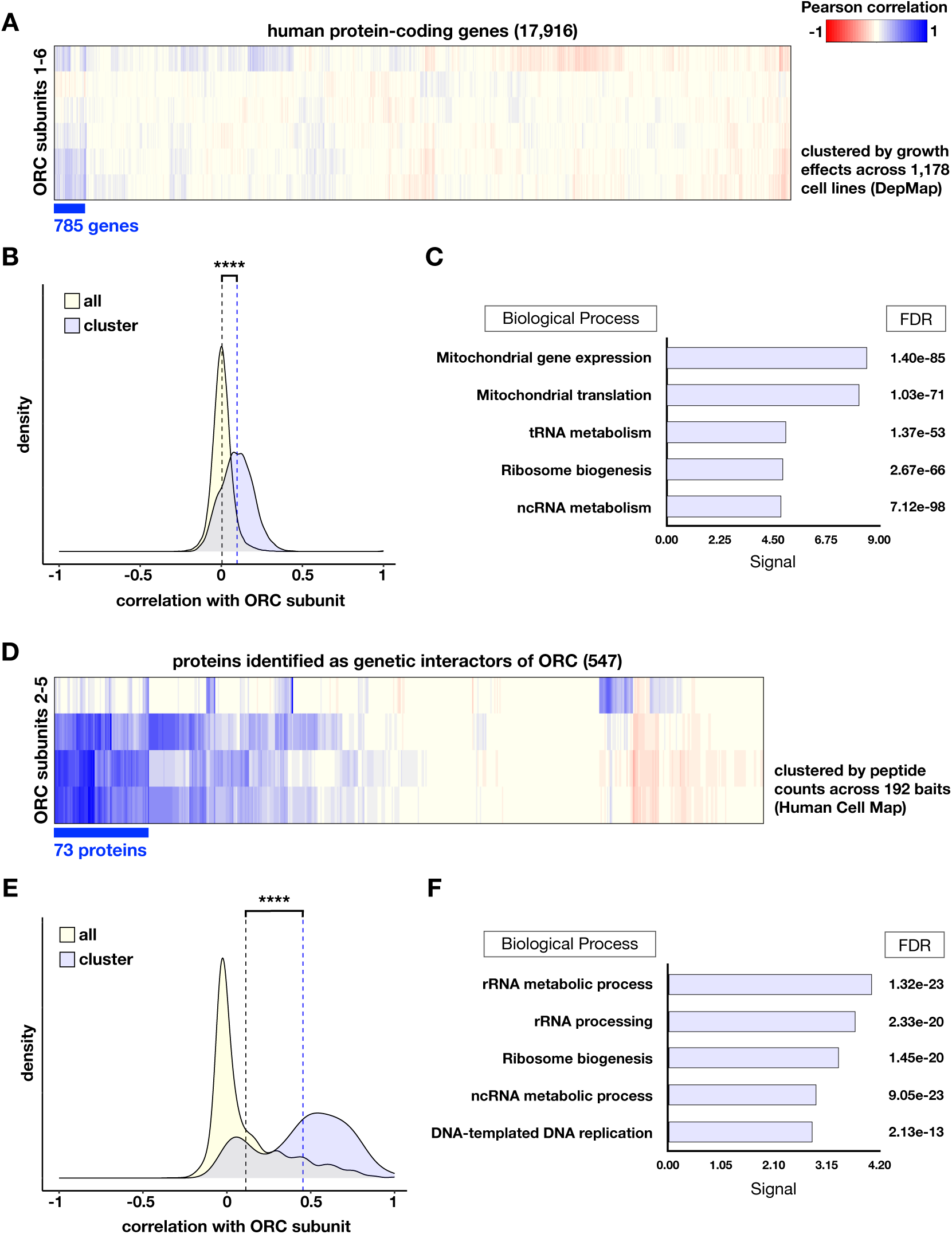
(A) Heatmap showing hierarchical clustering of human protein-coding genes by growth effect correlations with ORC subunits 1-6 using DepMap data. (B) Density plot of growth effect correlations of all genes available in DepMap with ORC1-6 versus only cluster genes with ORC1-6; dashed lines represent the mean ORC subunit correlation for each group, where black is the mean for “all” and blue is for “cluster”. (C) Gene ontology enrichment for genetic interaction cluster genes with corresponding False Discovery Rates (FDR). (D) Heatmap showing hierarchical clustering of genetic interactors by peptide count correlations with ORC subunits 2-5 using Human Cell Map data. (E) Density plot of growth effect correlations of all genetic interactors available in Human Cell Map with ORC2-5 versus only cluster genes with ORC2-5; dashed lines represent the mean ORC subunit correlation for each group, where black is the mean for “all” and blue is for “cluster”. (F) Gene ontology enrichment for dual-criteria (genetic and proteomic) ORC interactors with corresponding FDRs.

Genetic interactions often reflect functional dependencies that may or may not be mediated by molecular interactions between the protein products of two genes. Thus, I used molecular proximity in human cells as my second criterion to delineate genetic interactions with ORC that are more likely to stem from physical interactions. Using Human Cell Map data, I refined the ORC genetic interactome to a subcellular interactome by hierarchically clustering ORC genetic interactors by their peptide count correlations with ORC subunits [Fig.2D]. I note that Human Cell Map data is not available for all genetic interactors and ORC subunits, limiting this portion of the analysis to 547/785 genetic interactors and ORC subunits 2-5. Similar to the analysis of DepMap data, this identified a cluster of 73 proteins significantly enriched for positive peptide count correlations with ORC subunits (^r̅^ = 0.1131 for all, versus ^r̅^ = 0.4537 for cluster, p = 8.51e-83) [Fig.2E]. As expected, this subclustering step eliminated the likely indirect genetic interactors involved in mitochondrial gene expression, and further enriched the ORC interactome for factors involved in nuclear RNA metabolism, as well as DNA replication [Fig.2F]. This dual-criteria ORC interactome suggests that a subset of ORC proteins not only has functional dependencies on an array of RNA processes in the nucleus, but also cohabitates the subnuclear environments in which these processes occur.

Since both the DepMap and Human Cell Map datasets convolve all phases of the cell cycle and molecular pathways beyond genome replication, I next sought to identify an S phase- and replication-specific portion of the ORC interactome. To this end, using a physical association with DNA synthesis as my next criterion, I filtered the dual-criteria ORC interactome using the proteome identified by iPOND [Fig.3A]. As expected, using iPOND detection as a filtering criterion narrowed the ORC interactome to a group of 35 proteins significantly enriched for functions in DNA replication, but still containing many proteins with no previously described roles in genome replication [Fig.3B,C]. To focus my downstream analyses on novel candidates, I filtered the candidate pool using a curated list of established genome maintenance factors that have already been computationally screened for interactions with one another using a Structure Prediction and Omics-informed Classifier (SPOC) [Fig.3A].^39^ This yielded a final candidate list of 17 novel factors with the potential for replication-related ORC interactions during S phase. Unexpectedly, this novel candidate pool was significantly enriched for factors involved in the regulation of Pol III transcription and Pol III-derived RNA biogenesis [Fig.3B,D]. This confluence of genetic and proteomic interactions from three independent datasets suggests yet-to-be-defined molecular partnerships between ORC, Pol III-derived small ncRNA metabolism, and human genome replication.

**Figure 3.**
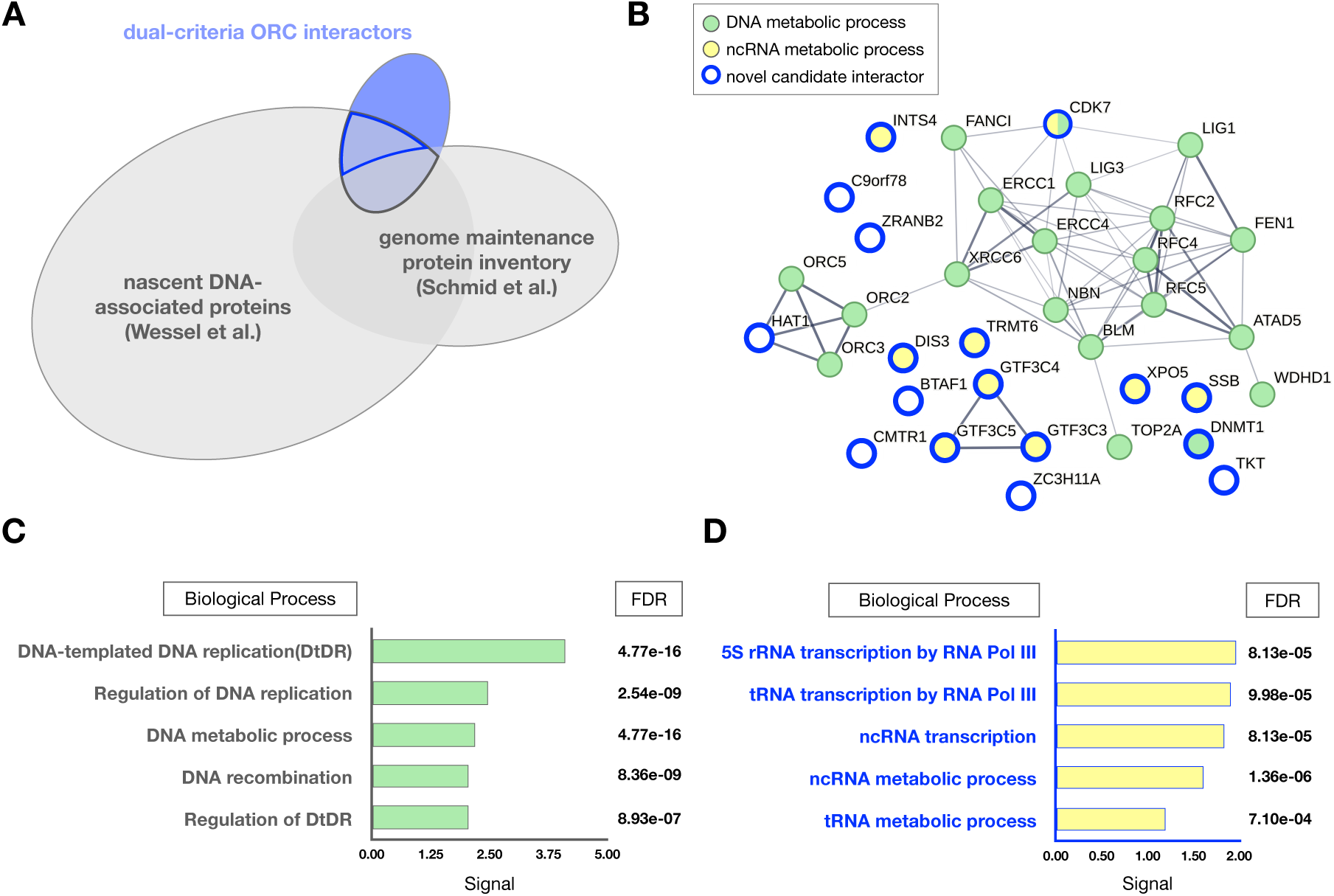
(A) Venn diagram showing the overlap between the dual-criteria ORC interactome, the nascent DNA-associated proteome identified by iPOND, and an inventory of canonical genome maintenance factors. (B) Proteins from the dual-criteria ORC interactome that were also identified by iPOND, represented as an interaction network predicted independently using the STRING database; edges and edge thickness represent data-supported interactions and interaction confidence, respectively. (C) Gene ontology enrichment for the iPOND-filtered ORC interactome with corresponding FDRs. (D) Gene ontology enrichment for proteins in the iPOND-filtered ORC interactome that are novel/not canonical genome maintenance factors, with corresponding FDRs.

### Structural prediction suggests direct interactions between ORC subunits and novel candidates involved in small noncoding RNA biogenesis, epigenetic regulation, and cell cycle control

Proteomic profiles obtained from proximity labeling and co-purification strategies can represent direct, indirect, stable, and transient physical associations. Aside from challenges in prioritizing potential protein interactions for follow-up, determining the precise molecular basis of numerous potential protein-protein interactions is often experimentally intractable. Increasingly, protein structure prediction offers tools to help gauge biomolecular interaction potential while minimizing experimental burden.^36,40^ To probe the possibility that the candidates I identified can engage in direct protein-protein interactions with ORC subunits, I leveraged the capacity of AlphaFold to perform multimeric structure prediction. AlphaFold uses predicted Template Modeling (pTM) scores as a heuristic for the overall quality of a predicted protein structure, taking into account per residue metrics like the Predicted Aligned Error (PAE). When two or more proteins are queried, this scoring is applied to predicted intermolecular contacts to generate an interface pTM (ipTM) score. As a confidence metric for intermolecular contacts, ipTM values offer a way to estimate the likelihood that two proteins interact, and thus can be used in a computational approximation of a protein-protein interaction screen.^39^ Given this logic, I used AlphaFold 3 (AF3) to screen for potential direct interactions between ORC subunits and novel candidates by generating three randomly seeded joint structure predictions for all possible ORC subunit-candidate protein (ORC-candidate) pairs and then determining their mean ipTM values [Fig.4A].

**Figure 4.**
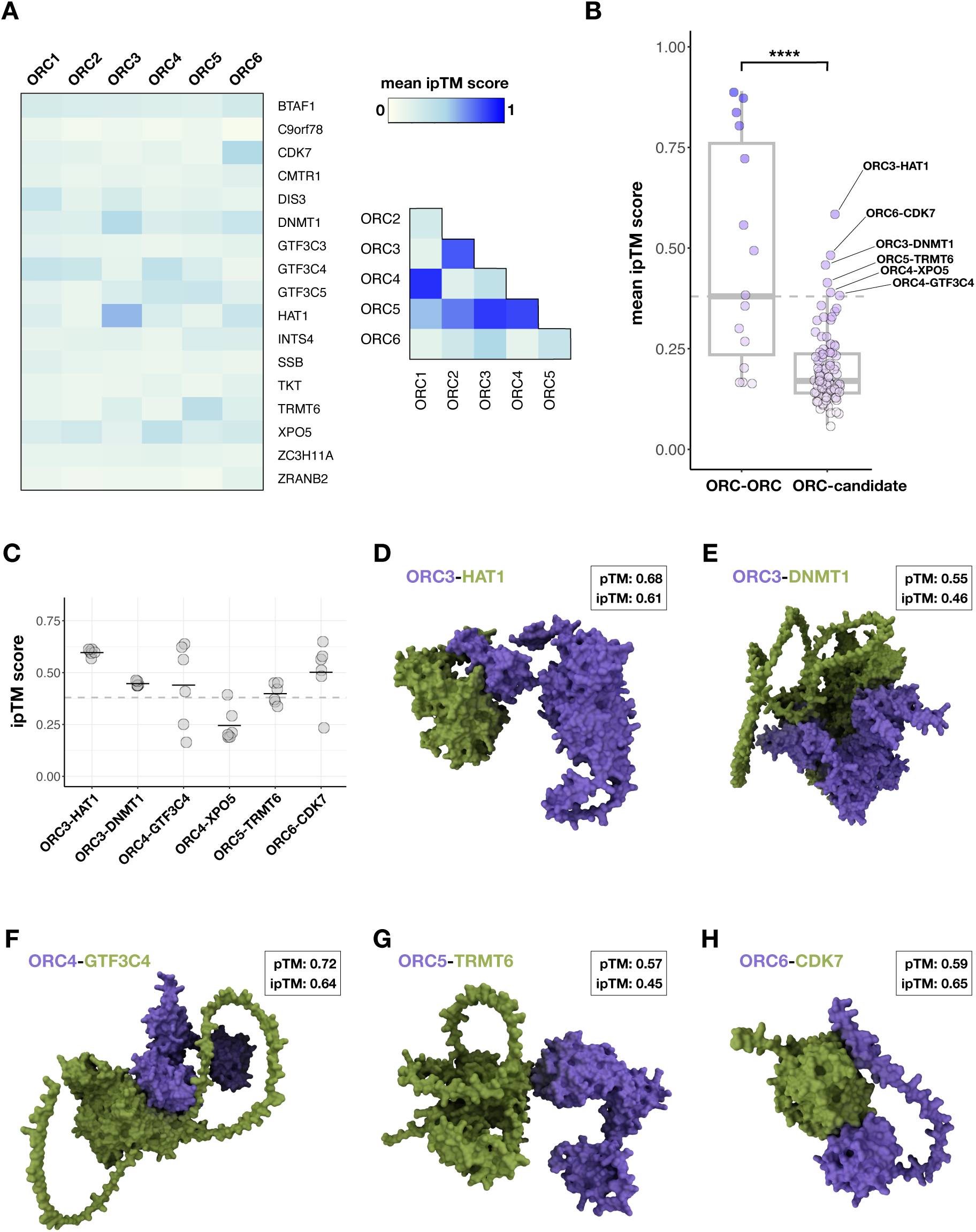
(A) Heatmaps showing mean ipTM scores for AF3 predictions querying ORC subunit-candidate protein pairs, and all heteromeric ORC subunit pairs. (B) Boxplots showing mean ipTM value distributions for heteromeric ORC subunit pairs (ORC-ORC) and ORC subunit-candidate protein pairs (ORC-candidate); the dashed line represents a cut-off of ≥ 0.38. (C) Jitter plots showing ipTM scores from retesting of top ORC-candidate pairs with additional AF3 predictions; black lines represent the mean. (D) 3D rendering of the top scoring predicted structure for an ORC3-HAT1 interface. (E) 3D rendering of the top scoring predicted structure for an ORC3-DNMT1 interface. (F) 3D rendering of the top scoring predicted structure for an ORC4-GTF3C4 interface. (G) 3D rendering of the top scoring predicted structure for an ORC5-TRMT6 interface. (H) 3D rendering of the top scoring predicted structure for an ORC6-CDK7 interface.

Consensus on meaningful ipTM values is still developing. Currently, general guidelines assert ipTM values lower than 0.6 suggest failed predictions, while some studies have had success pursuing predictions with ipTM values as low as 0.3.^41^ To address this ambiguity, I benchmarked mean ipTM values specifically for my purposes by also generating predictions for all heteromeric pairs of ORC subunits (ORC-ORC) [Fig.4A]; I reasoned this range of ipTM values could represent structurally well-characterized, physiological ORC subunit interactions, and in turn be informative for evaluating ORC-candidate ipTM scores. Comparison of the distributions of mean ipTM values for ORC-ORC and ORC-candidate pair predictions revealed the ORC-ORC scores were significantly higher on average than ORC-candidate scores (average mean ipTM score of 0.48 for ORC-ORC, versus 0.20 for ORC-candidate, p = 1.66e-5) [Fig.4B]. However, a subset of the ORC-candidate pairs were statistical outliers, with mean ipTM values of 0.38 or higher, placing them, coincidentally, at or above the median ORC-ORC value of 0.38. Thus, I chose a cut-off ipTM score of ≥ 0.38 to select predictions for follow-up, focusing on ORC-candidate pairs with mean ipTM scores that deviate substantially from the mean and are in the 50th percentile of ipTM scores for ORC self-associations. This identified 6 ORC-candidate pairs [Fig.4B].

Because AlphaFold predictions display stochasticity and sensitivity to initial state, I reran predictions for the 6 top hits and doubled the sampling to 6 randomly seeded predictions. Predictions for almost all top ORC-candidate pairs reproduced mean ipTM scores of 0.38 or higher, except ORC4-XPO5 [Fig.4C]. This result led to a final nomination of 5 proteins for hypothetical direct interactions with ORC subunits: HAT1, DNMT1, GTF3C4, TRMT6, and CDK7. I generated 3D renderings of predicted structures representing the highest ipTM value observed across predictions for each ORC subunit-candidate protein pair [Fig.4D-H]. These structures had pTM values ranging from 0.55-0.72, ipTM values ranging from 0.45-0.64, and no clashes reported. I note that I did not determine if these hypothetical protein-protein interfaces would interfere with interactions between ORC subunits; if experimentally validated, it will be important in future work to understand if these candidate interactions are mutually exclusive or compatible with formation of ORC subunit holocomplexes. In summary, the results of this structural prediction screen using AF3 further suggest that 5 of 17 candidate proteins nominated by genetic and proteomic interactions with ORC have the potential to directly bind ORC subunits.

### Structural prediction suggests direct interactions between ORC3 and Pol III-derived small noncoding RNAs

While some of the genetic and proteomic interactions with ORC subunits may be explained by protein-protein interactions, interactions with molecules other than proteins may also contribute to these interactomes. This is particularly relevant to the novel candidate pool, which is enriched for RNA-binding factors involved in the homeostasis of RNAs derived from Pol III transcription [Fig.3B,D]. Considering that multiple pathways in the novel candidate network converge on the homeostasis of 5S rRNA and tRNA, along with multiple reports that ORC can bind RNA^1,24^, I hypothesized that ORC subunits may have the ability to recognize these RNA species. To pursue this hypothesis, I leveraged AlphaFold’s recently developed ability to predict RNA structures.^36^ Combined with AF3’s capacity to do joint structure predictions, this creates exciting opportunities to predict novel ribonucleoprotein complexes.

Similar to my previous screen, I used AF3 to perform a joint protein-RNA structure prediction screen for plausible direct interactions between ORC subunits, 5S rRNA, and the methionyl tRNA (tRNA^met^) using mean ipTM scores; as a negative control, I also queried scrambled versions of both RNA sequences [Fig.5A]. Interestingly, the mean ipTM scores for ORC-RNA pair predictions were significantly higher on average than those for ORC-scramble RNA pairs (average mean ipTM score of 0.27 for ORC-RNA, versus 0.15 for ORC-scramble, p = 1.39e-2) [Fig.5B]. However, these differences are difficult to interpret without precedent for identifying meaningful ipTM scores for potential protein-RNA interactions. 5S rRNA and tRNAs have well-characterized protein interactions, and I exploited this knowledge to design positive and negative control test cases to evaluate the performance of AF3’s joint protein-RNA structure prediction and benchmark ipTM scores. To this end, I tested ribonucleoprotein complex predictions with known specific binders of 5S rRNA and tRNA^met^: RPL11 and RPL5 for 5S rRNA, and MARS1 and TRNT1 for tRNA^met^.^42–44^ Encouragingly, ipTM scores were high for predictions for known binders paired with their specific substrate, and low for pairings with non-specific substrates, including the scrambled versions of both 5S rRNA and tRNA^met^ [Fig.5A,B].

**Figure 5.**
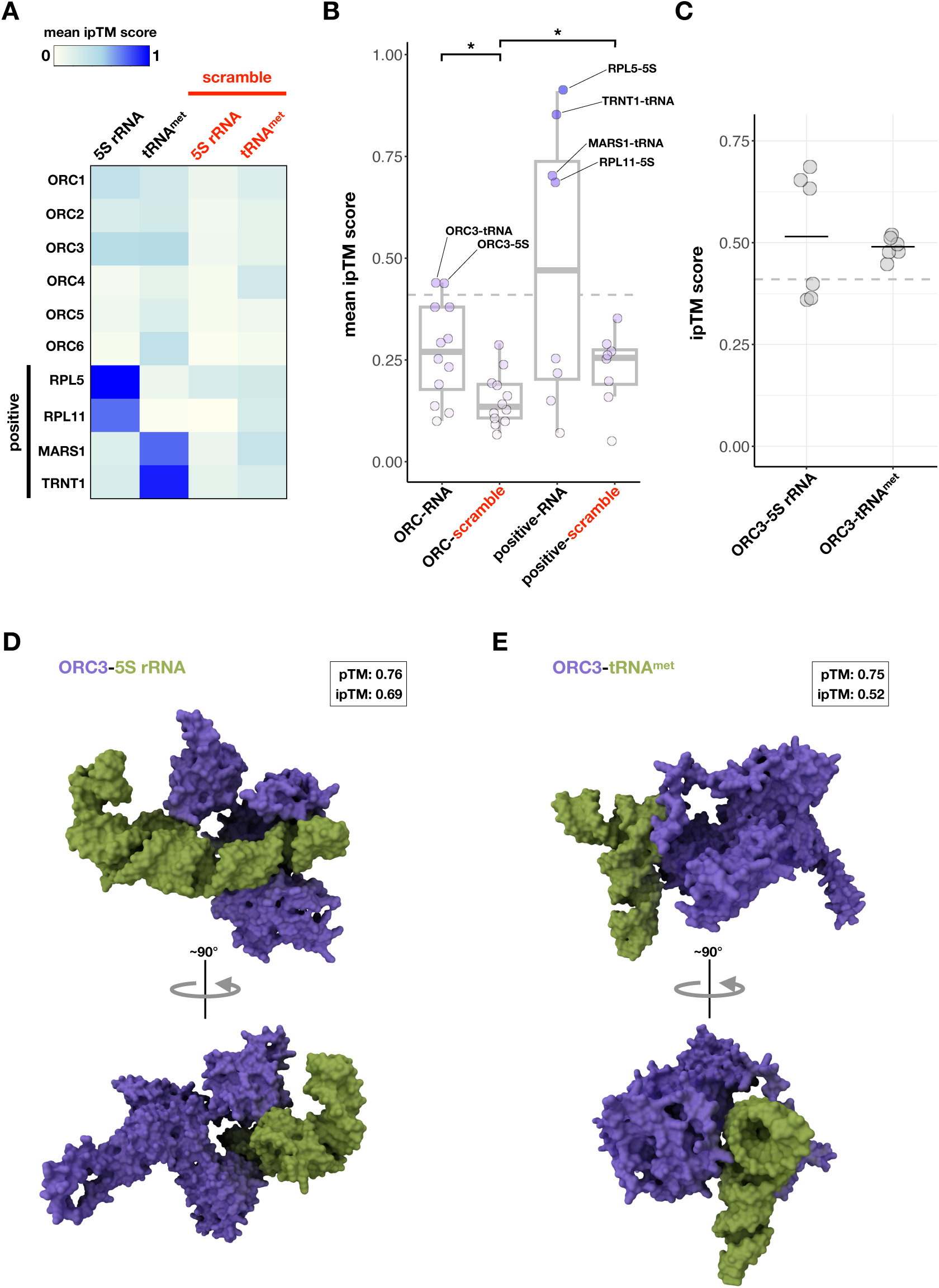
(A) Heatmap showing mean ipTM scores for AF3 predictions querying ORC subunits paired with 5S rRNA and tRNA^met^, along with positive control 5S rRNA- and tRNA-binding proteins. (B) Boxplots showing mean ipTM value distributions for ORC subunit-RNA pairs (ORC-RNA), ORC subunit-scramble RNA pairs (ORC-scramble), positive control protein-RNA pairs (positive-RNA), and positive control protein-scramble RNA pairs (positive-scramble); the dashed line represents a cut-off of ≥ 0.41. (C) Jitter plots showing ipTM scores from retesting of top ORC-RNA pairs with additional AF3 predictions; black lines represent the mean. (D) 3D rendering of the top scoring predicted structure for an ORC3-5S rRNA interface. (E) 3D rendering of the top scoring predicted structure for an ORC3-tRNA^met^ interface.

After confirming that AF3 joint protein-RNA structure predictions can produce ipTM scores in line with known biochemistry and biology, I reasoned that the ipTM scores for known RNA-binders paired with the scrambled negative control RNAs could be treated as a value range representing nonspecific interactions. Thus, I used the distribution of ipTM values generated from positive control protein-scrambled RNA pair predictions to establish an outlier-based cut-off of ≥ 0.41, enabling identification of ORC-RNA pair predictions with mean ipTM scores that deviate substantially from this background. This identified two ORC-RNA pairs: ORC3-5S rRNA, and ORC3-tRNA^met^ [Fig.5B]. I again performed 6 additional randomly seeded predictions for both pairs and confirmed the reproducibility of an average ipTM score of 0.41 or higher [Fig.5C]. I generated 3D renderings of predicted structures representing the highest ipTM value observed across predictions for each pair [Fig.5D,E]. The predicted ORC3-5S rRNA and ORC3-tRNA^met^ complexes had respective pTM values of 0.76 and 0.75, respective ipTM values of 0.69 and 0.52, and no clashes reported. I note, as with predictions for ORC subunit-candidate protein pairs, I did not investigate how these hypothetical ORC3-RNA interfaces might impact ORC3’s incorporation into an ORC protein holocomplex; this will also be an important goal for future study should these interactions be experimentally validated. To look for experimental evidence that ORC3 binds RNA in cells, I reviewed data generated by capture of RNA interactomes using click chemistry (RICK).^45^ RICK uses metabolic labeling of RNA to affinity-purify and proteomically profile crosslinked RNA-protein complexes from human cells. Excitingly, a high-confidence list of RNA-binding proteins identified by RICK contained ORC3, as well as 21 other factors in the dual-criteria, iPOND-filtered ORC interactome [Fig.S4]. Altogether, these results support hypotheses that ORC3 can form ribonucleoprotein complexes with small structured ncRNAs like 5S rRNA and tRNAs, and that protein-RNA interactions may contribute to the genetic and proteomic ORC interactomes identified earlier.

### Candidate ORC interactions reinforce and enhance a minimal model for origin activation

To conclude my investigation, I wondered if the results of my analyses could be phased with my minimal model for origin activation [Fig.1A]. Regarding hypothetical HAT-mediated feedback, my screen identified an obvious candidate that could serve this function: a potential interaction between ORC3 and the histone acetyltransferase HAT1 [Fig.6A]. Prevailing models argue that HAT1 acts primarily on free histone H4 during S phase, depositing the marks H4K5ac and H4K12ac.^46^ Evidence indicates these modifications are transient, facilitating chromatin reformation after replication and undergoing removal after histone incorporation. Additional evidence indicates more direct roles in DNA replication. HAT1 knockout in mouse embryonic fibroblasts impeded replication fork progression and induced DNA damage.^47^ In line with my findings, another study in yeast indicated that HAT1, while not required for replication initiation, physically and genetically interacts with yeast ORC.^48^ These reports align with my findings indicating that HAT1 genetically and proteomically interacts with ORC in human cells, possibly forming a complex with ORC3, and thus may have a role in the replication program.

**Figure 6.**
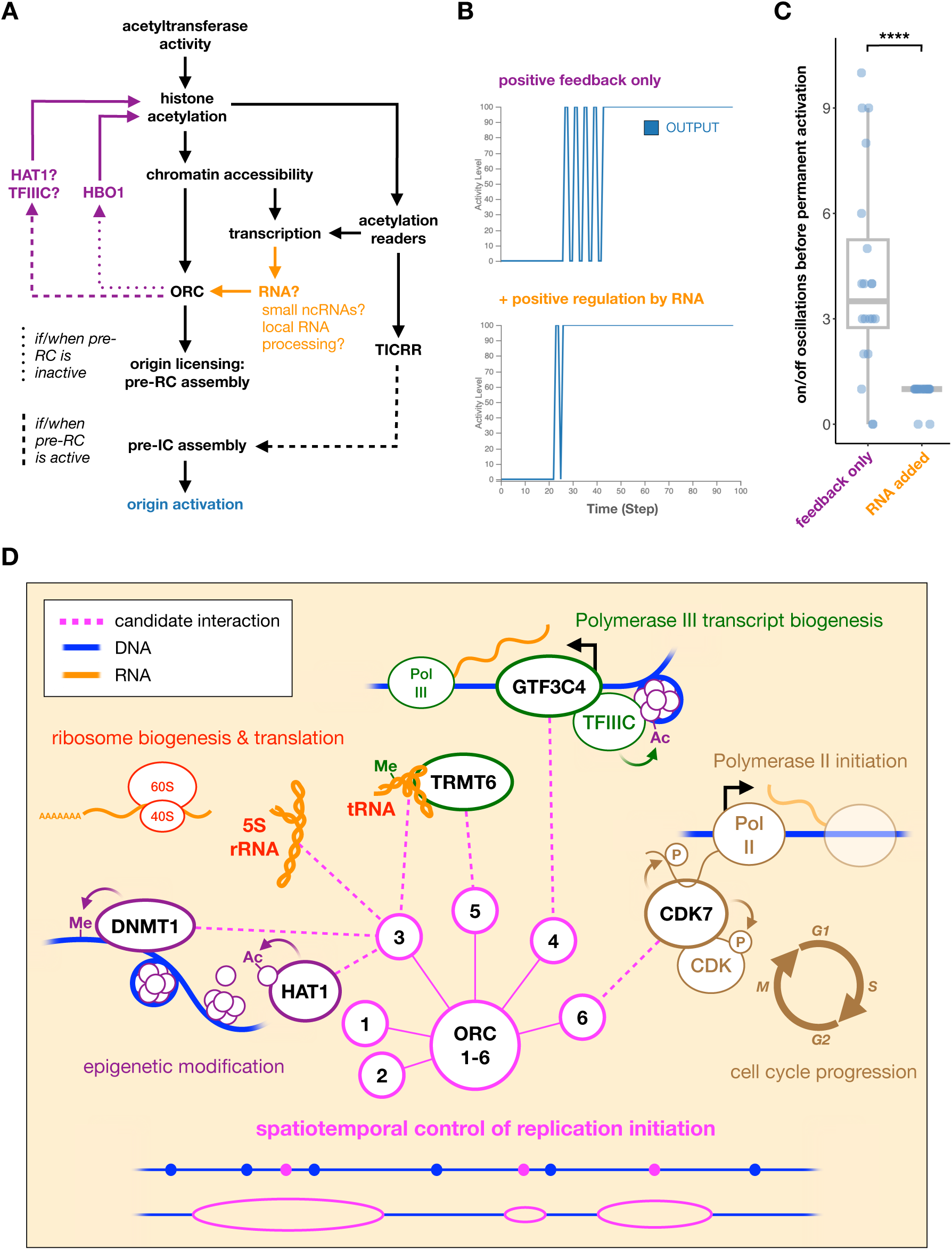
(A) Modified version of the model for origin activation from Fig.1A integrating chromatin acetylation, ORC, origin licensing, and transcription. (B) Outputs over time from simulations testing the model in A as Boolean logical models with: positive feedback representing ORC-mediated interactions with HBO1, HAT1, and/or TFIIIC (positive feedback only), and addition of a transcription-RNA-ORC path (+ positive regulation by RNA). (C) Boxplots showing quantification of output oscillations prior to permanent activation from 20 simulations, with and without addition of a transcription-RNA-ORC path (RNA added) to the model. (D) Diagram summarizing the hypothetical ORC interactions identified by this study, along with known protein functions, enzymatic activities, and relevant biological processes.

A less obvious candidate source of HAT activity is GTF3C4. Also known as TFIIIC90, GTF3C4 is a subunit of the TFIIIC complex. TFIIIC is part of the transcription factor ensemble that directly recognizes Type I and II promoters to recruit Pol III to 5S rRNA and tRNA genes.^49^ Interestingly, my analysis of genetic and proteomic interaction data identified three TFIIIC subunits: GTF3C3, GTF3C4, and GTF3C5. The TFIIIC complex has no previously described roles in genome replication. To the contrary, studies have suggested the Pol III machinery and its associated transcription factors are physical barriers to replication fork progression and cause stalling at Pol III transcription sites.^50^ However, my analysis infers that subunits of the TFIIIC complex are co-essential and cohabitate with ORC subunits, and suggests that GTF3C4 can form a complex with ORC4. Interestingly, GTF3C4 and the TFIIIC complex possess HAT activity, with multiple studies indicating that TFIIIC has the ability to acetylate histone H3 both in vitro and in cells.^51–53^ If the ORC3-HAT1 and ORC4-GTF3C4 interactions posited by my work can be experimentally validated, it would be interesting to probe the possibility that ORC interactions can congregate these proteins and origins after licensing and during S phase to modulate H3 and H4 acetylation, and in turn origin activation.

Given my model is histone acetylation-centric, there is also opportunity to incorporate transcription and RNA biogenesis into the network. Multiple studies have provided evidence that ORC binds RNA in vitro and in cells, and have suggested that ORC-RNA interactions link transcription and replication patterning.^24,54–56^ My results suggest that ORC function is linked to the biogenesis and local processing of RNA, particularly in relation to Pol III transcription. My analysis of genetic and proteomic data links ORC and replication to the TFIIIC complex, La protein (SSB), which binds nascent Pol III-derived RNAs, and the tRNA methyltransferase cofactor TRMT6. Furthermore, my structural prediction results inform hypotheses that ORC5 directly interacts with TRMT6 and ORC3 directly interacts with Pol III-derived small ncRNAs. Motivated by these findings, I sought to test the impact of introducing a transcription-RNA-ORC regulatory axis into my model. To this end, I created additional paths in my logical model by which open chromatin and acetylation readers promote transcription, transcription promotes RNA, and RNA promotes ORC activity [Fig.6A]. Simulated outputs from this augmented model appeared to more efficiently enter a permanently active state than the model with positive feedback alone [Fig.6B]. To measure this effect, I quantified the number of on/off oscillations prior to permanent activation across 20 simulations for both models. Quantification confirmed that the additional positive regulation of ORC by a transcription-RNA pathway significantly decreased anticipatory oscillations (mean number of anticipatory oscillations = 4.2 with positive feedback only, versus 0.9 with a transcription-RNA path added, p = 1.22e-5), further promoting a switch-like output [Fig.6C]. In total, these results inform a model in which ORC interactions with HATs and local RNA biogenesis are self-reinforcing, promoting accumulations of ORC and histone acetylation at transcriptionally active nuclear domains for efficient origin licensing and activation.

Lastly, my data mining and structural predictions also identified potential ORC subunit interactions with the DNA methyltransferase DNMT1 and the cyclin-dependent kinase CDK7. DNMT1 is best known for propagating cytosine methylation patterns onto newly synthesized DNA.^57^ DNMT1 carries out this function through recruitment to nascent DNA and recognition of hemimethylated CpG dinucleotides. While DNMT1’s primary role is to propagate pre-existing methylation patterns, evidence suggests DNMT1 performs de novo methylation at appreciable rates.^58^ Though DNMT1’s activity can be considered replication-coupled, there is minimal evidence that it is physically coupled to the replication process; some studies have suggested a direct but inessential interaction with the processivity clamp PCNA.^59,60^ My analysis reveals signatures of functional and physical association between DNMT1 and ORC, and posits a direct interaction between DNMT1 and ORC3. If experimentally confirmed, it would be interesting to explore how an ORC-DNMT1 interaction could help target DNMT1 to replication initiation sites to facilitate prompt association with nascent, hemimethylated DNA, or, conversely, modulate DNMT1’s activity to preserve the methylation status of efficient, early origins, which tend to be in hypomethylated CpG island promoters.^1^

CDK7 is a multifunctional cyclin-dependent kinase, functioning in both transcriptional regulation and cell cycle progression. CDK7 regulates RNA polymerase II (Pol II) by phosphorylating Pol II’s carboxy-terminal domain, promoting the transition of promoter-paused Pol II complexes into elongation.^61^ Additionally, CDK7 can phosphorylate and activate other CDK proteins that further promote transcription and also cell cycle progression. Interestingly, CDK7 inhibition has been shown to induce genome instability and replication stress.^62^ My analyses suggest CDK7 is a genetic and proteomic interactor of ORC, with potential to form a complex with ORC6. ORC6 is unique amongst ORC subunits, with reports that it is variably associated with ORC protein holocomplexes, potentially dispensable for replication initiation in some contexts, and involved in genome surveillance in S phase.^63,64^ Regardless, if an ORC6-CDK7 interaction can be validated, it will be interesting to determine if CDK7 phosphorylation influences origin activity, or if CDK7 phosphorylates ORC subunits. Additionally, the potential for a CDK7-ORC interaction to link the regulation of Pol II transcription and replication patterning would complement my minimal model for origin activation, bolstering the network with additional interactions connecting ORC and transcription.

In summary, this work integrates process, proximity, and prediction to infer a network of hypothetical interactions with human ORC proteins that may have functions beyond origin licensing [Fig.6D]. All of the hypothetical physical interactions put forth by this study require validation by genetic, molecular, and/or biochemical experiments. The results of follow-up experiments will be important in determining if the approach I have employed here is an effective way to accelerate discovery in molecular biology by minimizing the number of experiments required to derive tenable hypotheses and models. If confirmed, these interactions, broadly, have the potential to integrate replication patterning in new ways with epigenome maintenance, Pol III transcript biogenesis, regulation of Pol II transcription, ribosome biogenesis and translation, and cell cycle control [Fig.6D]. By extension, these findings also have potential implications for genome stability. At a systems level, my results suggest that ORC proteins operate at an interface of genome maintenance and nuclear RNA metabolism, or across these domains. Numerous studies have observed that defects in nuclear RNA metabolism (transcription, RNA processing and packaging, etc.) lead to replication stress and genome instability. Prevailing models for this instability invoke excessive hybridizations of nascent RNA to DNA, termed R-loops, and increases in transcription-replication conflicts as causes of DNA damage.^65–67^ However, my previous work demonstrated that many reports of excess R-loop formation were confounded by reliance on the nonspecific, RNA-binding S9.6 antibody in imaging applications, and may have misinterpreted abnormal nuclear accumulations and distributions of RNA as changes in R-loops.^68^ This suggests that abnormalities in the abundance and organization of RNA in the nucleus consistently coincide with replication stress and genome instability. There is mounting evidence that DNA and RNA metabolism have reciprocal relations, with RNA biogenesis contributing to DNA repair and canonical DNA repair factors acting in RNA biogenesis.^69,70^ These findings favor models centered on genome maintenance factors having beneficial physical interactions with RNA and RNA biogenesis under steady-state conditions, and suggest that defective RNA homeostasis could perturb genome maintenance by means other than R-loop formation and transcription-replication conflicts. My findings suggest that ORC function, and thus the replication program, may be governed by such principles.

## Supplemental data

**Figure S1.**
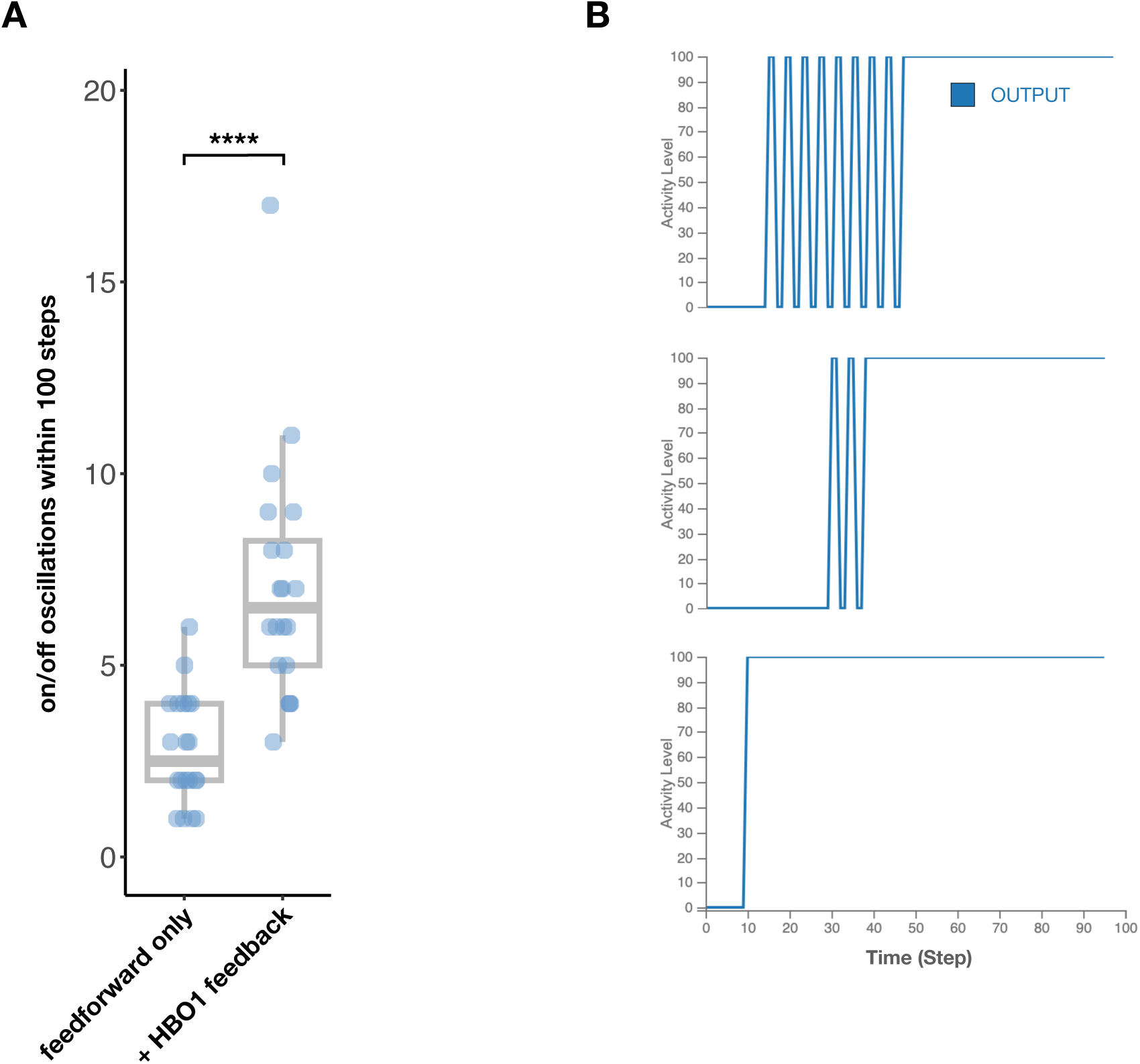
(A) Boxplots showing the number on/off oscillations within 100 steps from 20 simulations of the feedforward model in Fig.1A with and without the addition HBO1-mediated positive feedback. (B) Plots of bistable output over time from 3 different simulations of the model in Fig.1A with both HBO1- and hypothetical HAT-mediated feedback incorporated.

**Figure S2.**
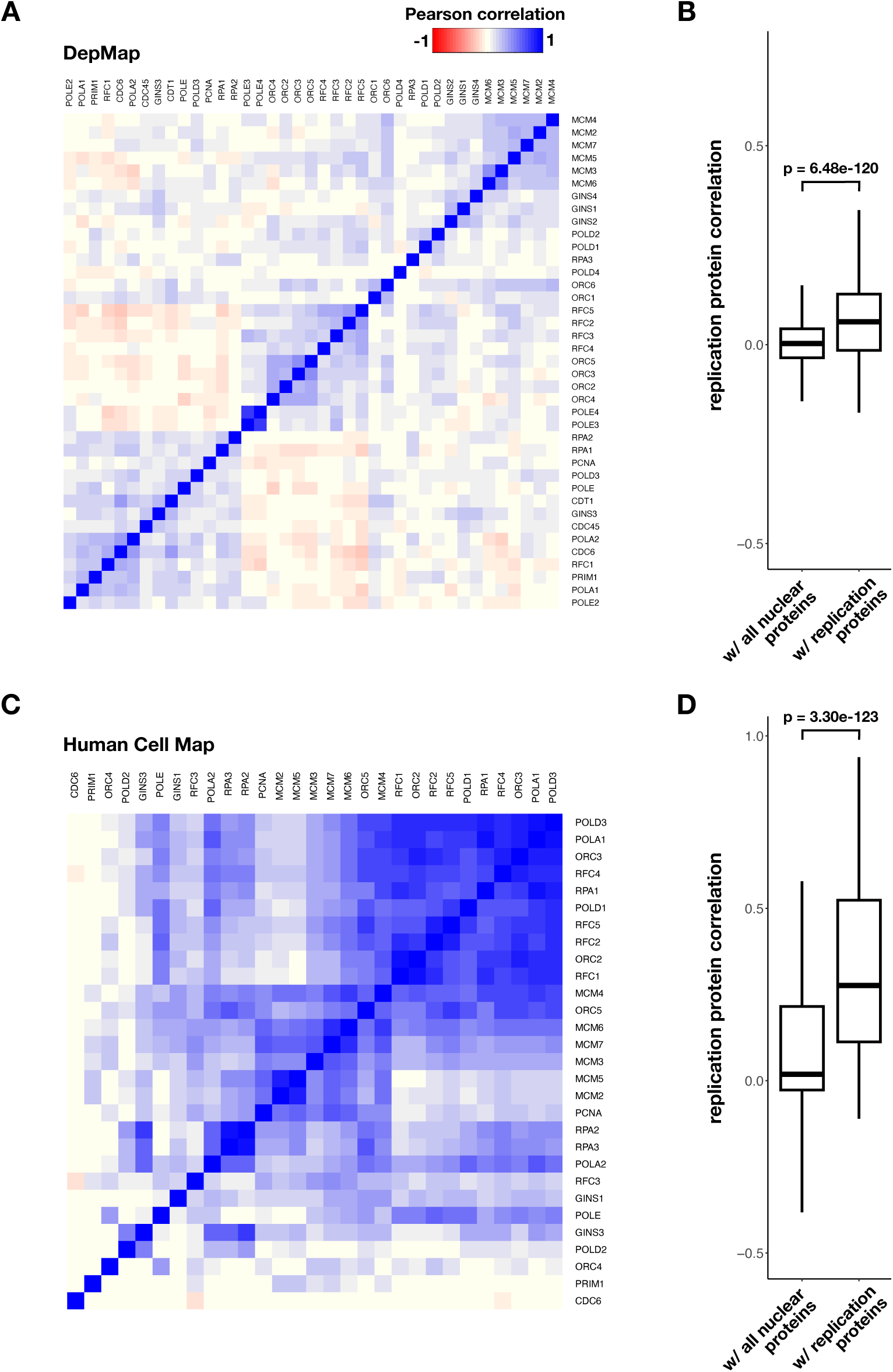
(A) Heatmap showing an all-by-all matrix of core replication proteins hierarchically clustered by growth effect data from DepMap. (B) Boxplots comparing the distributions of growth effect correlations of replication proteins with all available nuclear proteins versus replication proteins with replication proteins; self-correlations were excluded. (C) Heatmap showing an all-by-all matrix of core replication proteins hierarchically clustered by proteomic data from the Human Cell Map project. (D) Boxplots comparing the distributions of peptide count correlations of replication proteins with all available nuclear proteins versus replication proteins with replication proteins; self-correlations were excluded.

**Figure S3.**
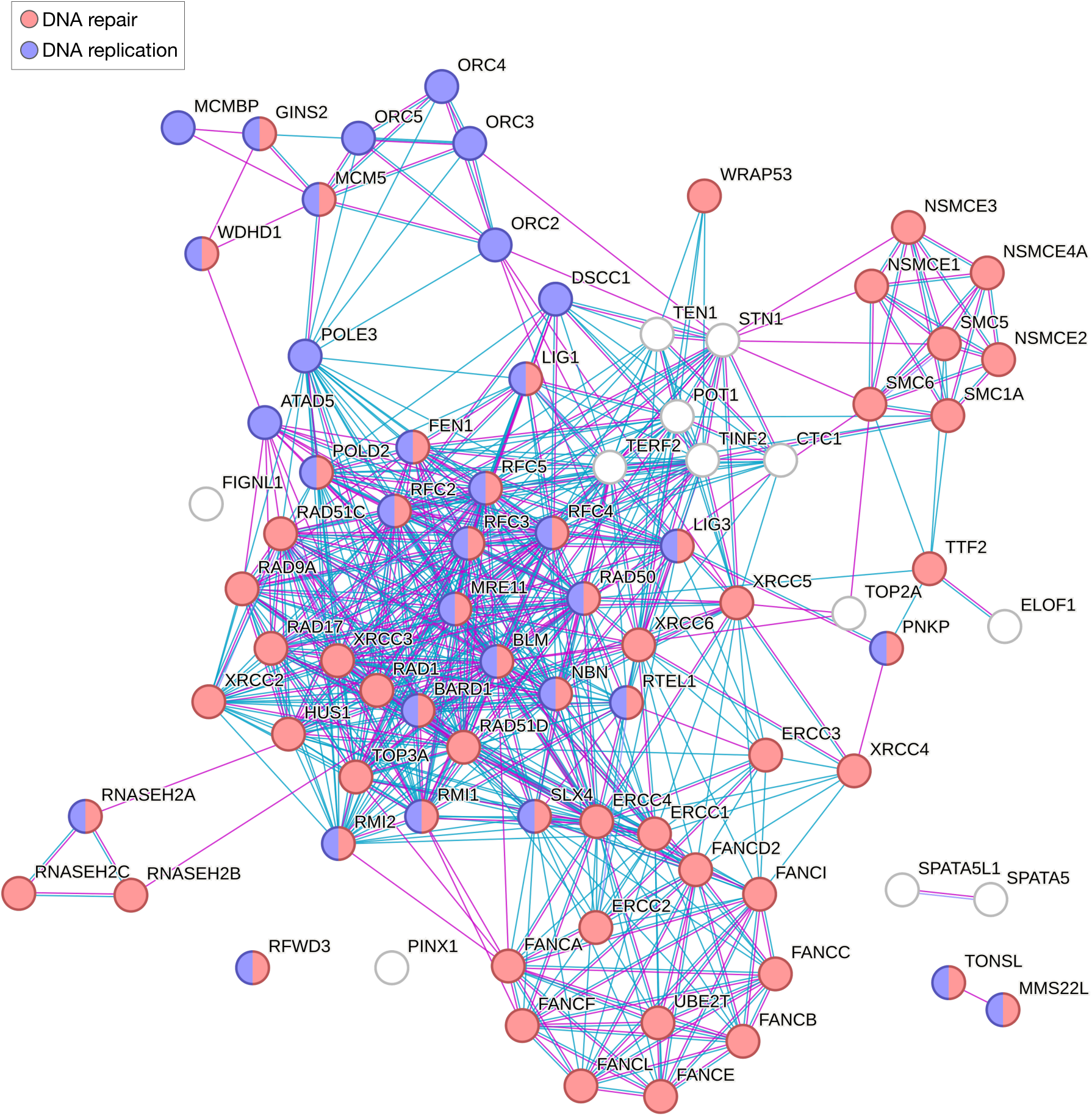
Proteins from the ORC genetic interactome with established roles in genome maintenance^39^, represented as an interaction network generated using the STRING database; edges represent interaction evidence.

**Figure S4.**
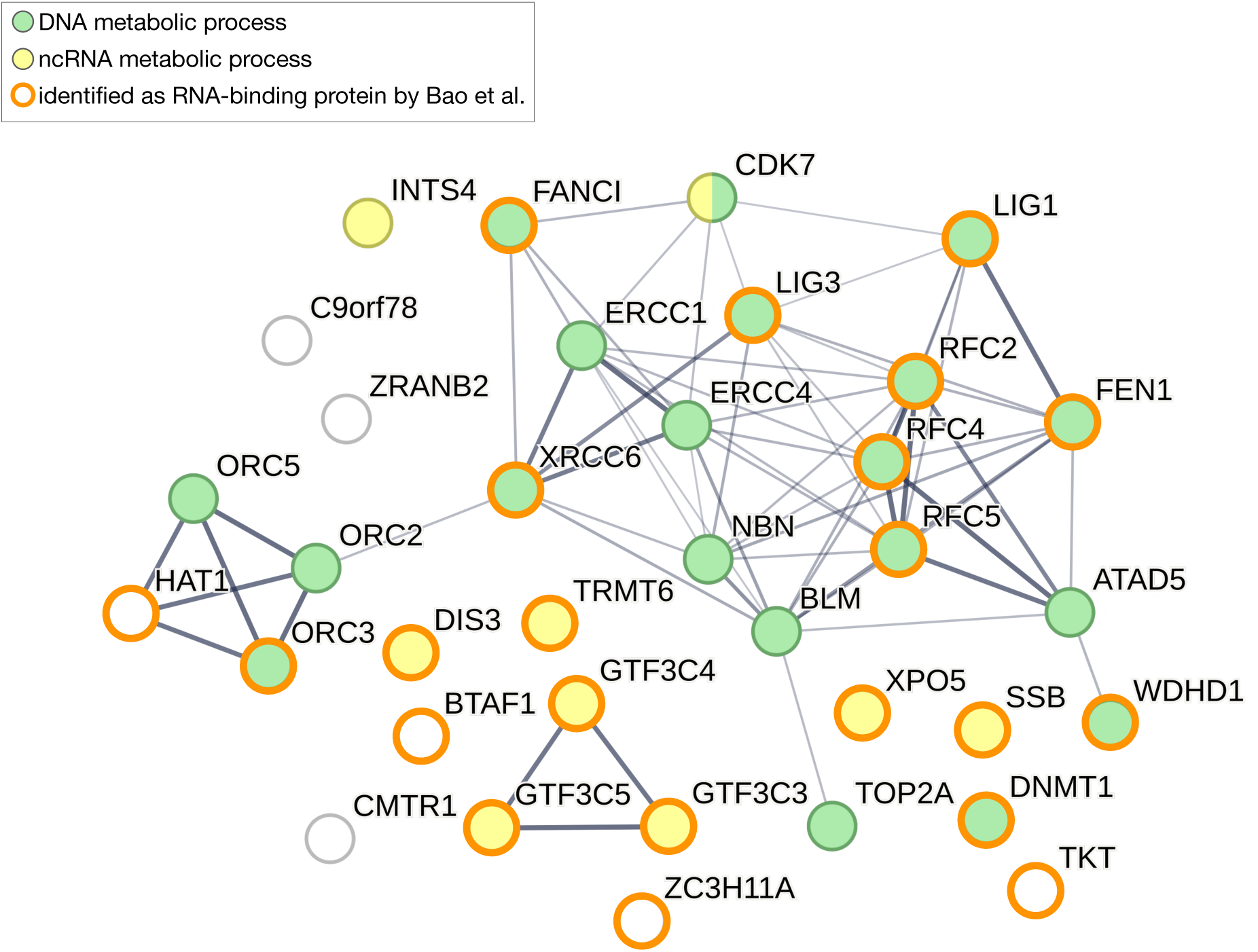
Proteins from the dual-criteria, iPOND-filtered ORC interactome that have been identified as RNA-binding proteins by Bao et al.^45^, shown using the interaction network from Fig.3B.

